# The pace of mitochondrial molecular evolution varies with seasonal migration distance

**DOI:** 10.1101/2023.08.01.551470

**Authors:** Teresa M. Pegan, Jacob S. Berv, Eric R. Gulson-Castillo, Abigail A. Kimmitt, Benjamin M. Winger

## Abstract

Animals that engage in long-distance seasonal migration experience strong selective pressures on their metabolic performance and life history, with potential consequences for molecular evolution. Species with slow life histories typically show lower rates of synonymous substitution (dS) than “fast” species. Previous work has suggested that long-distance seasonal migrants have a slower life history strategy than short-distance migrants, raising the possibility that rates of molecular evolution may covary with migration distance. Additionally, long-distance migrants may face strong selection on metabolically important mitochondrial genes owing to their long-distance flights. Using over 1000 mitochondrial genomes, we assessed the relationship between migration distance and mitochondrial molecular evolution in 39 boreal-breeding migratory bird species. We show that migration distance correlates negatively with dS, suggesting that the slow life history associated with long-distance migration is reflected in rates of molecular evolution. Mitochondrial genes in every study species exhibited evidence of purifying selection, but the strength of selection was greater in short-distance migrants, contrary to our predictions. This result may indicate selection for cold tolerance on mitochondrial evolution among species overwintering at high latitudes. Our study demonstrates that the pervasive correlation between life history and molecular evolutionary rates exists in the context of differential adaptations to seasonality.

## Introduction

Species’ traits are the product of their genome and their environment, but in turn, traits and the environment also shape the molecular evolution of the genome. For example, metabolically-demanding traits influence molecular evolution of mitochondrial genes (e.g. Shen et al. 2009; Chong and Mueller 2013; Strohm et al. 2015). More broadly, traits associated with the slow-fast continuum of life history (Stearns 1983) are correlated with rates of molecular evolution (Bromham 2020) such that life history evolution is thought to alter the pace of a lineage’s molecular clock (Hwang and Green 2004; Moorjani et al. 2016). Environmental pressures associated with seasonality can influence life history (Varpe 2017) and metabolic demands (Weber 2009; Chen et al. 2018), suggesting that variation in adaptation to seasonality could have molecular evolutionary consequences. However, the linkages between molecular evolution and differential adaptations to seasonality are rarely explored.

In this study, we investigate how patterns of mitochondrial molecular evolution are related to variation in seasonal migration distance. Migratory animals survive harsh seasonal conditions on their breeding grounds by temporarily departing until conditions improve (Winger et al. 2019). Migration distance varies across species, ranging from short-distance movements within an ecoregion to hemisphere-crossing journeys. Long-distance seasonal migration requires high metabolic performance (Weber 2009), with potential implications for the dynamics of selection on the metabolically-important mitochondrial genes (Shen et al. 2009; Strohm et al. 2015). Migration distance has also been recognized as an important axis of life history variation (the balance between annual survival and reproduction) in birds (Greenberg 1980; Møller 2007; Bruderer and Salewski 2009; Winger and Pegan 2021). Through effects on life history (Bromham 2020), migration distance may therefore also influence molecular evolutionary rates, but this relationship has not been directly assessed. Here, we assess how migration distance correlates with mitochondrial molecular evolution within the community of migratory birds breeding in the highly seasonal North American boreal region, and we explore the roles of life history and metabolic adaptation in mediating a relationship between molecular evolution and seasonal migration.

### Metabolic adaptation, life history, and mitochondrial molecular evolution

Reliance on locomotion (migration) for adaptation to seasonality may influence selection on mitochondrial genes, which play an important role in metabolism. Mitochondria typically experience purifying selection (i.e. selection that reduces genetic variation) because most mutations in these genes are deleterious to fitness (Nei et al. 2010; Nabholz et al. 2013; Popadin et al. 2013). Prior studies have shown that purifying selection tends to be stronger in the mitochondria of mobile animal species compared with less-mobile relatives. This pattern has been demonstrated in comparisons between flighted and flightless birds (Shen et al. 2009) and insects (Mitterboeck et al. 2017; Chang et al. 2020), between migratory and nonmigratory fishes (Strohm et al. 2015), and between amphibians (Chong and Mueller 2013) and mollusks (Sun et al. 2017) with different locomotory modes. Within flighted birds, species with slow flight and those that rely on soaring (versus flapping) have been shown to experience relaxed mitochondrial purifying selection compared with faster-flying species (Shen et al. 2009; De Panis et al. 2021). These studies suggest that mitochondrial genotype plays an especially important role in fitness for organisms that rely on high-energy locomotion, including migratory birds. Metabolic demand may be strongest in long-distance migrating species if these demands primarily arise from locomotion. However, species that migrate short distances within seasonal high latitude regions may require alternative metabolic adaptations for dealing with harsh seasonal conditions since their shorter migrations do not allow them to fully escape cold, resource depleted winters (Winger et al. 2019). The effect of variation in seasonal migration distance on mitochondrial purifying selection is unknown.

A second and distinct way in which seasonal migration may influence molecular evolution is through its effect on life history and, consequently, molecular evolutionary rate. The slow-fast continuum of life history is commonly characterized by “life history traits” that underly or correlate with differing rates of growth, survival, and reproduction (Read and Harvey 1989; White et al. 2022). Within major lineages of plants, bacteria, vertebrates, and invertebrates, species with “slow” life history (i.e., long generation time, low annual fecundity, large size; Stearns 1983) also exhibit slower molecular substitution rate than “fast” species (i.e., shorter generation time, higher annual fecundity, and smaller size; Nabholz et al. 2008a; Smith and Donoghue 2008; Thomas et al. 2010; Weller and Wu 2015). Within migratory birds breeding in the temperate zone, seasonal migration distance covaries with annual fecundity and survival such that long-distance migrants show “slower” life history (i.e., higher annual survival, lower annual fecundity) than short-distance migrants (Greenberg 1980; Bruderer and Salewski 2009; Winger and Pegan 2021). As such, variation in migration distance across species may have consequences for molecular evolutionary rates. Specifically, the synonymous substitution rate “dS” is often correlated with the slow-fast life history continuum (Nikolaev et al. 2007, Bromham et al. 2015, Hua et al. 2015; Table 1). Prior studies suggest that life history may influence dS through effects on DNA replication rate or selection for mutation avoidance (reviewed in Bromham 2020), because dS is thought to primarily reflect the underlying mutation rate when synonymous mutations are selectively neutral (Kimura 1983; Nei et al. 2010; Lanfear et al. 2014). Direct estimates of nuclear germline mutation rates support the hypothesis that species-level variation in mutation rate correlates with life history traits (Bergeron et al. 2023).

**Table 1.** Definitions of abbreviations for molecular substitution rates and population genetic parameters and predictions for their relationships with migration distance.

| Concept | Abbr. | Description and assumptions | Predictions (this study) |
| --- | --- | --- | --- |
| Synonymous substitution rate | dS | Assuming synonymous sites evolve neutrally, dS primarily reflects $\mu$ (Nei et al. 2010; Lanfear et al. 2014) | Negative relationship between migration distance and dS |
| Nonsynonymous substitution rate | dN | Assuming nonsynonymous sites are generally deleterious, dN is influenced by both $\mu$ and $N_e$ (reviewed in Nei 2005) | NA |
| dN/dS ratio | dN/dS | Assuming nonsynonymous mutations are generally deleterious, dN/dS reflects strength of purifying selection on dN while accounting for variation in $\mu$. Low dN/dS = strong purifying selection. (Nei 2005; Kryazhimskiy and Plotkin 2008) | Negative relationship between $\theta$ and dN/dS, reflecting the influence of $N_e$ on dN/dS.<br>Negative relationship between migration distance and dN/dS, indicating positive relationship between migration distance and purifying selection strength. |
| Mutation rate | $\mu$ | May be influenced by life history; reviewed in (Bromham 2020) | NA, $\mu$ not measurable in our data |
| Effective population size | $N_e$ | Defined as the ideal population size experiencing the same level of genetic drift as observed in the data (Waples 2022). Estimated in mitochondrial data as $\theta / \mu$. (Watterson 1975; Nabholz et al. 2008a) | NA, see $\theta$ |
| Theta | $\theta$ | Population genetic parameter representing genetic variation. Assuming low variation in $\mu$, variation in $\theta$ primarily reflects variation in $N_e$ | Negative relation between $\theta$ and dN/dS and between $\theta$ and $\pi N / \pi S$ |
| Synonymous nucleotide diversity | $\pi S$ | Population genetic parameter representing population-level nucleotide diversity at synonymous sites. | NA |
| Nonsynonymous nucleotide diversity | $\pi N$ | Population genetic parameter representing population-level nucleotide diversity at synonymous sites. | NA |
| $\pi N / \pi S$ ratio | $\pi N / \pi S$ | Reduction of $\pi N$ compared to $\pi S$ is expected to reflect natural selection, but the relationship is more complex than in dN/dS | Negative relationship between migration distance and $\pi N / \pi S$, indicating positive relationship between migration distance and selection. Negative relationship between $\theta$ and $\pi N / \pi S$, indicating purifying selection on nonsynonymous polymorphisms. |

### Predicting the relationship between seasonal migration distance and molecular evolution

Long-distance migratory birds have been shown to exhibit a slower life history than sympatric breeding short-distance migrants (Winger and Pegan 2021, Fig. 1). Thus, long-distance migrants travel farther in each migratory trip than short-distance migrants and may also require more trips per lifetime to achieve the level of lifetime fitness of short-distance migrants (Møller 2007). Owing to the metabolic demands of migration and the importance of repeated migration success for fitness in long-distance migrants, the migratory phenotypes of these species are thought to be under strong variation-reducing natural selection (Conklin et al. 2017). As such, we hypothesize that long-distance migrants exhibit both lower dS (which could reflect selection against mutation in the mitochondria; Hua et al. 2015) and stronger purifying selection in their mitochondrial genes than short-distance migrants.

**Figure 1.**
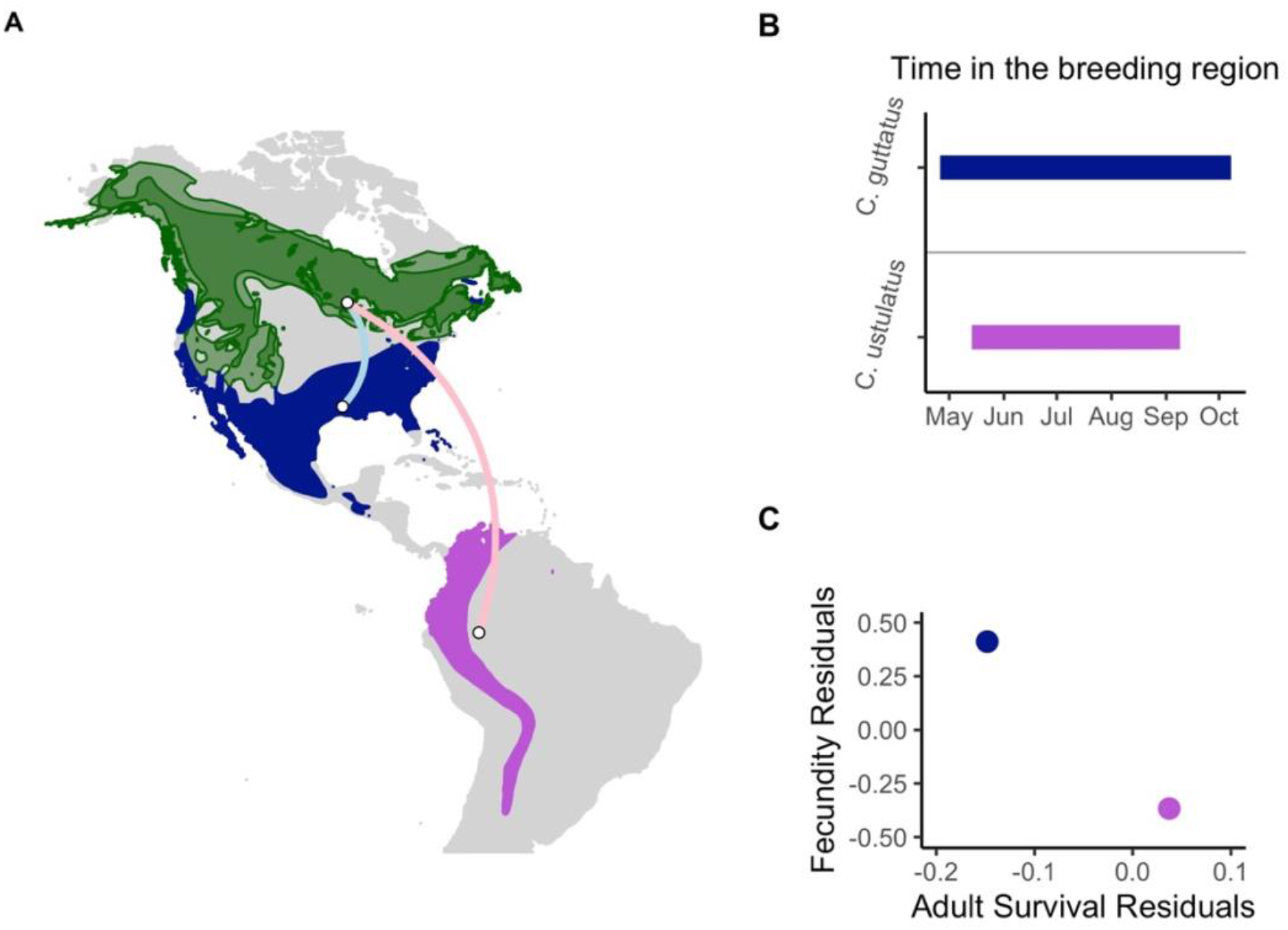
An example contrast between a shorter-distance migrant *Catharus guttatus* and a closely related longer-distance migrant *Catharus ustulatus swainsoni* illustrates the relationship between migration distance and life history in our study system. Both species have broadly overlapping breeding ranges (green), but *C. guttatus* (dark blue nonbreeding range) migrates a shorter distance (blue migratory route) than *C. u. swainsoni* (purple nonbreeding range, pink migratory route) (panel A). Accordingly, *C. guttatus* spends more time in its breeding range than *C. u. swainsoni* (panel B). With more time in the breeding range and the possibility of raising a second brood, the short-distance migrant has higher fecundity but lower adult survival—i.e., faster life history—than the long-distance migrant (panel C, showing model residuals from mass-corrected analysis of fecundity and survival). The short-distance migrant spends the winter in colder, more resource-depleted regions than the long-distance migrant. Figure and data adapted from Winger and Pegan (2021). Our sampling for this study occurred only within the eastern boreal belt (Fig. S1).

To test these hypotheses, we examined the relationship between migration distance and rates of molecular evolution of the mitochondrial coding genes in a community of small-bodied migratory songbirds breeding in the boreal forests of North America. The 39 co-distributed species we studied are ideal for investigating the effects of migration distance on molecular evolution because they vary greatly in migration distance (e.g., Fig. 1, Table S1), yet they otherwise share similar breeding habitat, population history, and body mass (Winger and Pegan 2021). This system allows us to test hypotheses about migration distance while minimizing variation in other traits that could influence molecular evolution. We assessed effects of migration distance on dS (synonymous substitution rate) and dN/dS (purifying selection) in a phylogenetic framework (Lartillot and Poujol 2011) with full mitochondrial gene sets we sequenced for 39 species. Further, we used population genetic datasets from all mitochondrial genes that we generated for 30 of the species (for a total of 1008 samples used across all analyses) to assess effects of migration distance on purifying selection within populations (πN/πS) and to account for the potentially confounding influence of effective population size (N_e_) on molecular evolution.

## Methods

### Study system

We focused on 39 species of migratory birds breeding in the North American boreal forest, representing 11 families (Table S1). These are the same species for which a correlation between migration distance and the slow-fast life history continuum, independent of body size, was demonstrated using data on annual fecundity and survivorship (Winger and Pegan 2021). We focus our analyses on co-distributed populations of the eastern boreal belt of North America (Omernik 1987, Fig. S1). Some species’ breeding ranges extend into other ecoregions (e.g., the mountain west or the temperate forests south of the boreal zone), but in these cases we only analyze samples from the boreal portion of the range so as to assess sympatric populations. The species in the dataset exhibit broad variation in migration distance, with their geographic range centroids shifting between 1048 km and 7600 km between the breeding and non-breeding periods (Fig. 2, Table S1; Winger and Pegan 2021). These centroid shifts represent migratory strategies ranging from short-distance movements within the temperate region to the movement of an entire population across ocean and land barriers from North America to South America. All species are less than 100 g in mass (range of mean mass across species is 6-87 grams; Table S1) and are broadly similar in habitat use. They are all territorial species with socially monogamous breeding systems, which suggests that they probably do not vary substantially in population sex ratio (which can affect N_e_), although empirical sex ratio data is not available for these species. Small songbirds are typically capable of breeding in their second year, and this is true of all species in our study that have been assessed (Billerman et al. 2022). Additionally, the species share relatively similar demographic histories, with population expansions estimated to have mostly occurred during the period of glacial retreat that preceded the Last Glacial Maximum (∼57,000 years before present; Kimmitt et al. 2023).

**Figure 2.**
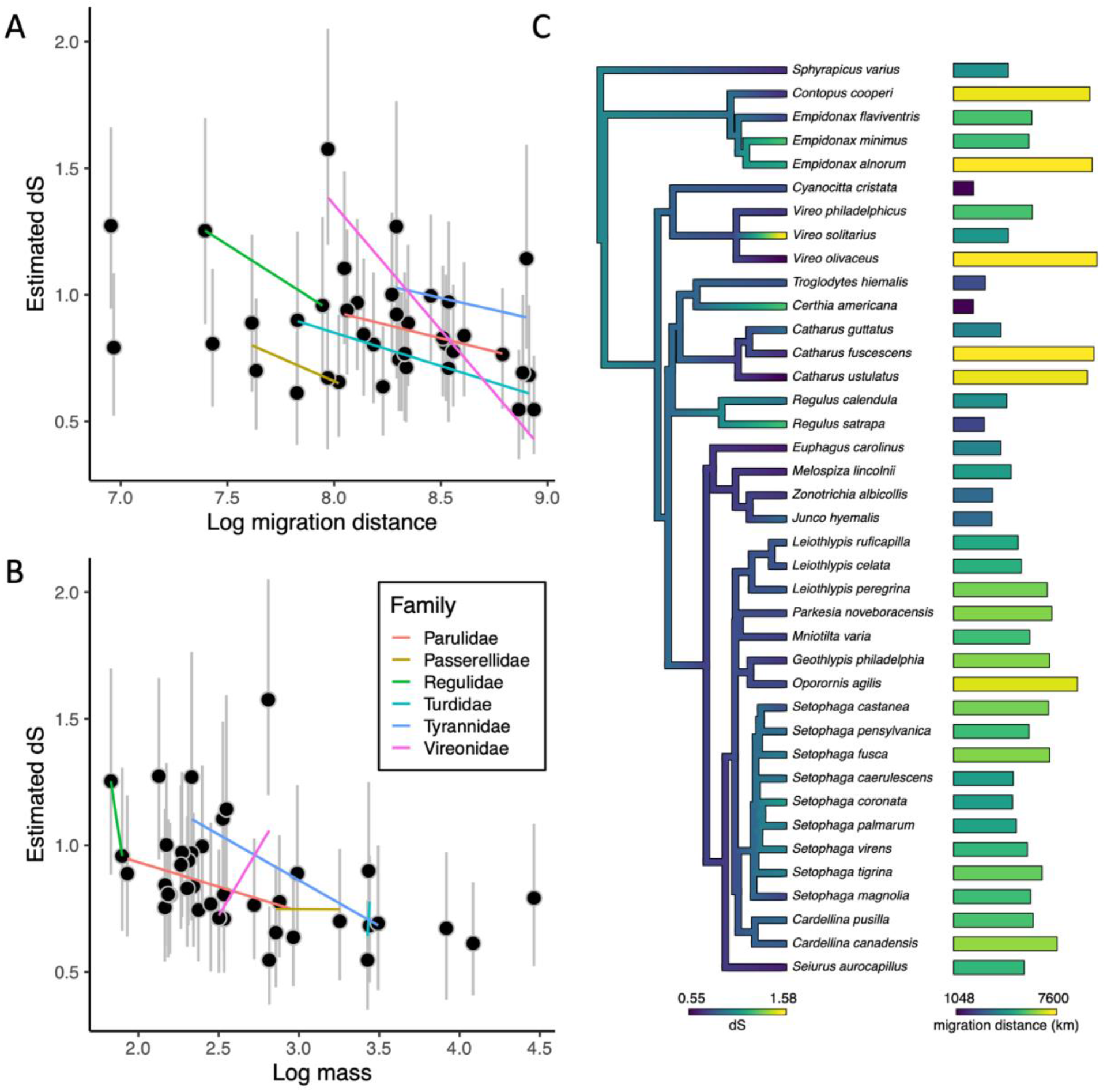
dS vs. traits associated with life history (A, B) and a phylogenetic tree showing dS and migration distance for each species (C). In panels A and B, posterior mean tip estimates of dS (black dots) from Coevol are shown compared to migration distance (A), and mass (B) from models using our full species set. Gray vertical bars indicate 95% credible intervals for each estimate. These analyses reveal that both migration distance and mass have a negative relationship with dS. Plotted lines use linear models to visualize the relationship between estimated tip dS and a given covariate within each family of birds (when represented in our dataset by two or more species), demonstrating a consistently negative relationship between dS and migration distance within and among major clades in our system. In panel C, the phylogenetic tree was created in phytools (Revell 2012) and is colored based on posterior mean tip and node estimates of dS from Coevol.

### Life history covariates: Migration distance and mass

Direct measurements of migration distance of individuals are lacking for most of the species in our system, so we used the distance between the centroid of a species’ breeding range and the centroid of its nonbreeding range to represent the migration distance of the species. Although the distance between centroids does not represent individual variation in migration distance within a species, this metric captures broad differences in migratory strategies between species. Our method for calculating the distance between range centroids is described in detail in Winger and Pegan (2021). We included mass as a covariate in our analyses because body mass and rates of molecular evolution are often associated (Figuet et al. 2014; Nabholz et al. 2016), and the relationships between survival and fecundity and migration distance demonstrated by Winger and Pegan (2021) were recovered after controlling for the effect of mass. We obtained mass data from Dunning (2008) and Billerman et al. (2022).

### Sampling and DNA sequencing

Our analysis of the relationship between migration distance and dS requires one mitochondrial genome for each species in the study, while analyses of N_e_ and πN/πS require population-level sampling. For our analysis of dS, we obtained whole mitochondrial genomes from one individual of each of the 39 species in our study by sequencing DNA as described below. Samples were collected during the breeding season from near the longitudinal center of the boreal forest (Manitoba, Minnesota, or Michigan; S2). For two species (*Contopus cooperi* and *Euphagus carolinus*), we sequenced DNA from individuals sampled during migration in Michigan from collision mortalities.

For our population-level analyses, we generated a large dataset of 999 additional mitochondrial genomes for 30 of the 39 species, building on a dataset of 19 species from Kimmitt et al. (2023). Our larger dataset includes complete coding sequences for 8 to 49 individuals per species (mean 33 individuals per species; Table S1). These individuals were sampled during the breeding season across a longitudinal transect of the boreal forest from Alberta to the northeastern United States (Fig. S1, Table S2). Except for 24 blood samples from New York state, all sequences we used came from frozen or ethanol-preserved tissue samples provided by several museum institutions (Table S2; *Acknowledgments*).

We obtained high-depth mitochondrial genomes captured as a byproduct from low-coverage whole genome sequencing, as described in detail in Kimmitt et al. (2023). Briefly, sequencing libraries were prepared using a modified Illumina Nextera library preparation protocol (Schweizer et al. 2021) and sequenced on HiSeq or NovaSeq machines using services provided by Novogene and the University of Michigan Advanced Genomics Core. We used NOVOPlasty v4.3.1 (Dierckxsens et al. 2016) to assemble mitochondrial contigs, specifying a target genome size of 20-30 kb and using a k-mer of 21. We provided NOVOPlasty with a conspecific mitochondrial seed sequence (Table S1) for each species. We annotated the contigs built by NOVOPlasty using Geneious Prime 2020.2.2 (https://www.geneious.com) with copies of mitochondrial genes from GenBank (Table S1). Whenever applicable in the filtering and analysis steps described below, we used options specifying the vertebrate mitochondrial code.

Our initial dataset across all species contained mitochondrial sequences from 1229 total individuals. To ensure data quality, we used BLAST (https://blast.ncbi.nlm.nih.gov/Blast.cgi) to check species identity and we removed samples with evidence of species misidentification, chimerism, or introgression from related species (14 samples removed). We aligned and translated sequences with the R package DECIPHER v2.18.1 (Wright 2016) and we visually inspected each alignment, ensuring that sequences contained no premature stop codons or other alignment issues. We used DECIPHER to remove partial stop codons and to remove the untranslated C in the ND3 sequence of woodpecker (Picidae) species (Mindell et al. 1998). As our population analyses require complete data matrices, we excluded individuals with incomplete datasets (those with assemblies that were missing genes and/or with ambiguous base calls; 202 samples removed) and we concatenated the 13 genes for each remaining individual. An additional 5 individuals were removed during populational structure analysis, described below. This data filtering resulted in 1008 complete mitochondrial coding sequences: 999 individuals across 30 species used in the population genomic analyses plus one sequence for each of the 9 additional species we used only in the interspecific Coevol analyses. The full list of samples, including those that were removed from the analyses, can be found in Table S2.

### Accounting for effects of N_e_ on substitution rates using θ

Many parameters of molecular evolution are fundamentally associated with effective population size (N_e_), so estimating N_e_ provides important context for our analyses. Variation in N_e_ can cause variation in substitution rates because in theory, the efficiency of natural selection in purging deleterious mutations is determined by the balance between strength of selection and strength of drift, which is reflected by N_e_ (Ohta 1992). Specifically, studies on empirical populations have demonstrated that populations with small N_e_ typically show weaker purifying selection (i.e., higher dN/dS, e.g., Popadin et al. 2007, Leroy et al. 2021; and higher πN/πS, e.g., Chen et al 2017). Similarly, nearly neutral theory suggests that N_e_ can influence dS when synonymous sites are not selectively neutral (as in Chamary et al. 2006). That is, under nearly-neutral theory, synonymous sites with weak influence on fitness may be under purifying selection in populations with large N_e_ and not in populations with small N_e_, which could result in lower dS for species with high N_e_. Several recent studies find correlations between traits associated with life history and genetic diversity, suggesting that species with “slow” life histories often have low N_e_ (Romiguier et al. 2014; Brüniche-Olsen et al. 2021; De Kort et al. 2021). For these reasons, when assessing the relationship between seasonal migration and molecular evolution, we tested whether molecular rate variation across species could alternatively be explained by confounding variation in N_e_.

We used genetic diversity (θ) as a proxy for N_e_. Effective population size (N_e_) can be calculated based on θ and mutation rate (Watterson 1975, Nabholz et al. 2008b; Table 1), but accurate estimates of mitochondrial mutation rate are lacking for most non-model organisms. Accordingly, many empirical studies interested in N_e_ focus on genetic diversity, which is thought to reflect the harmonic mean of N_e_ over time and which does not require mutation rate information to calculate (e.g., Ellegren and Galtier 2016; Hague and Routman 2016). We hereafter use the genetic diversity parameter θ as a proxy for N_e_. We used LAMARC v2.1.10 (Kuhner 2006) to estimate θ for each species. We imported our sequence data into LAMARC after converting our concatenated fasta files into the phylip format for each species. We used the program’s likelihood-based method in 10 initial chains (samples = 500, discard = 1000, interval = 20) and 2 final chains (samples = 10,000, discard = 1000, interval = 20). We used the F84 model of molecular evolution and we provided a separate transition/transversion ratio for each species using values we calculated from population sequence datasets using the R package ‘spider’ (Brown et al. 2012). We examined the output for each species to check for chain convergence and we ran two replicate chains for each species to make sure they produced consistent results. For 5 species (*Leiothlypis ruficapilla, Setophaga castanea, Setophaga coronata, Setophaga fusca,* and *Vireo olivaceus*), we repeated LAMARC for 25 initial chains instead of 10 to improve convergence and used the values from these longer runs.

### Population Structure

Our population-level analyses (estimation of θ and πN/πS) assume that there is no geographic population genetic structure within the samples used. To check this assumption, we calculated mitochondrial genetic distance between all individuals within each species using “nei.dist()” from the R package poppr v2.9.3 (Kamvar et al. 2014) and created a neighbor-joining tree with “nj()” from the R package ape v5.6-2 (Paradis and Schleip 2019). We identified and removed 4 individuals from *Regulus satrapa* and one individual from *Oporornis agilis,* all from Alberta in the far western part of our sampling area, that were clearly genetically distinct from all other samples in their respective species. Otherwise, there was little evidence of geographic genetic structure in the mitochondrial genome in these species.

### Estimating dS and dN/dS and their correlations with traits associated with life history

We used Coevol v1.6 (Lartillot and Poujol 2011) to evaluate associations between migration distance and molecular evolutionary rates using a single representative of each species. Coevol uses a Bayesian phylogenetic framework to estimate dS and dN/dS and to simultaneously measure the relationship between these traits and covariates of interest (migration distance, mass, and θ). We included mass to account for the expected relationship between mass and molecular rates (Nabholz et al. 2016). Models with mass also provide a useful point of comparison, allowing us to ask whether migration distance correlates with dS and dN/dS to the same extent as (or more or less than) this well-studied life history trait. Similarly, including θ in the models allows us to assess whether variation in N_e_ accounts for differences in molecular evolutionary rates.

We provided Coevol with one complete mitochondrial coding sequence from each species and a phylogenetic tree (Fig. 2) we built with data from birdtree.org (Jetz et al. 2012) as described in Pegan and Winger (2020). In brief, we sampled 2000 trees comprising all North American bird species from the Jetz et al. dataset and we used the python package “DendroPy” (Sukumaran and Holder 2010) to generate a consensus tree. We then trimmed this tree to include only the 39 species used in this study. Importantly, Coevol only uses the phylogenetic tree for topological information, and the program estimates branch lengths from the separate sequence data during modeling (Lartillot and Poujol 2021). Coevol does not require information about mutation rates. Because Coevol does not take topological uncertainty into account, we investigated potential effects of phylogenetic tree topology on our results by sampling 10 random marginal trees from the original tree dataset (trimmed to include only relevant species) and running a Coevol model on each, which we found to produce consistent results (Table S3).

We created two data subsets for Coevol models: one subset contained all species in the study and included mass and migration distance as covariates. The other subset included the 30 species for which we had population-level data available; for these we included θ as a covariate in addition to mass and migration distance. For each data subset, we ran Coevol in 4 chains: two replicate chains with the option “dnds” (estimating dS; models 1 and 2, Table 2) and two with “dsom” (estimating dN/dS; models 3 and 4, Table 2). We let each chain run for approximately 20000 steps and examined the resulting trace files to ensure convergence and evaluate estimated sample sizes (ESS). All models converged and all parameters had ESS > 300. We removed the first 500 steps of each chain and thinned the chain to retain every 10^th^ step to reduce autocorrelation. Replicate chains produced highly similar estimates, and the values we report here represent the mean value of estimates made by each replicate chain. Full Coevol model output for each chain is presented in Tables S4 and S5.

**Table 2.**
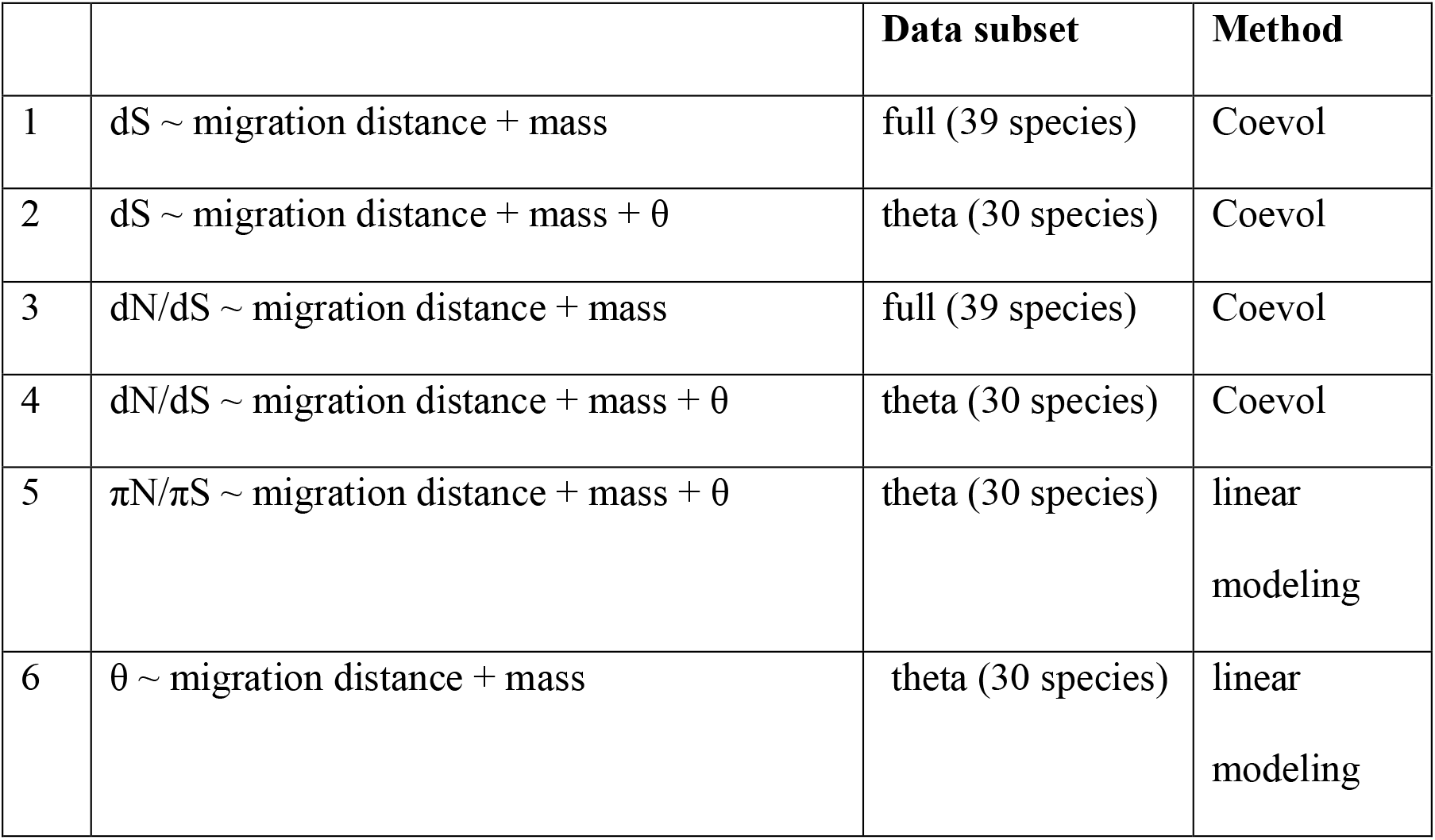
A summary of analyses. Models 1 and 2 use Coevol test our hypothesis that synonymous substitution rate (dS) is influenced by migration distance, with mass and θ (model 2 only) as additional covariates. Models 3 and 4 use the same approach with Coevol to estimate correlations between traits of interest and dN/dS. Models including θ use only 30 species because we did not have population-level data available to estimate θ for all 39 species. Coevol does not analyze molecular evolutionary parameters based on population-level data, so we used linear modeling to test whether traits of interest influence πN/πS (model 5). Finally, we also used linear modeling to test for potential confounding relationships between θ and life history-associated traits of interest (mass and migration distance; model 6).

|  |  | Data subset | Method |
| --- | --- | --- | --- |
| 1 | dS ~ migration distance + mass | full (39 species) | Coevol |
| 2 | dS ~ migration distance + mass + $\theta$ | theta (30 species) | Coevol |
| 3 | dN/dS ~ migration distance + mass | full (39 species) | Coevol |
| 4 | dN/dS ~ migration distance + mass + $\theta$ | theta (30 species) | Coevol |
| 5 | $\pi N/\pi S$ ~ migration distance + mass + $\theta$ | theta (30 species) | linear<br>modeling |
| 6 | $\theta$ ~ migration distance + mass | theta (30 species) | linear<br>modeling |

The method implemented in the Coevol software provides correlation coefficients between substitution rates and each covariate, as well as partial correlation coefficients (which hold constant the effects of other covariates in the model). Each correlation or partial correlation coefficient is accompanied by a posterior probability. In the case of Coevol, posterior probabilities near 0 indicate strong support for a negative relationship, while posterior probabilities near 1 indicate strong support for a positive relationship (Lartillot and Poujol 2021). *πN/πS*

πN/πS is a population genetic summary statistic representing the amount of nonsynonymous vs synonymous polymorphism within a population. This value is measured by comparing individuals within a species rather than by comparing between species in a phylogenetic framework (and thus cannot be estimated by Coevol). We estimated πN/πS from each species with population-level fasta alignments, using the python package egglib v3.1.0 (De Mita and Siol 2012) to create a “CodingDiversity” class with attributes describing the number of codons with synonymous or nonsynonymous polymorphisms. Predictions about the effect of purifying selection on polymorphisms are more complex than predictions about substitution rates because within-population variation can be purged by strong directional selective sweeps in addition to purifying selection (Kryazhimskiy and Plotkin 2008). Nevertheless, we predict a negative relationship between migration distance and the πN/πS ratio, which could result from either stronger purifying or stronger positive selection in long-distance migrants on mitochondrial function. In either case, such a relationship would broadly support the hypothesis that migration distance covaries with the dynamics of selection on the mitochondrial genome. We used linear modeling to test for an effect of migration distance, mass and θ on πN/πS (Tables S6, S7). Prior to linear modeling, we centered and standardized our predictors using the function “standardize” from the R package “robustHD” (Alfons 2021) with the mean value of each predictor as the center. We used a similar linear modeling approach to test whether θ exhibits a relationship with mass or migration distance to ensure that apparent relationships between these traits and molecular rates are not confounded by correlation with θ.

For each response variable (θ and πN/πS; Tables S6, S7), we first created a model with all covariates of interest. We then used the function “phylosig()” from the R package phytools v0.7-70 (Revell 2010) to test for phylogenetic signal in the model’s residuals (Revell 2012). For both response variables, the estimate of lambda (phylogenetic signal) was low (< 0.2) and the p-value for evidence of phylogenetic signal was > 0.8, so we proceeded with linear modeling rather than using models with phylogenetic covariance matrices. For each response variable, we created a null (intercept-only) model with no predictors and models with all possible combinations of our predictors of interest, and we used the function “model.sel()” from the R package MuMIn v1.43.17 (Bartón 2019) to compare the models’ AICc.

## Results

For each model, we report correlation coefficients between traits of interest (migration distance, mass, or θ) and molecular evolutionary rates (dS or dN/dS). We assess the strength of evidence for correlations using posterior probabilities (*pp*), which are close to 0 in the case of a strong negative correlation and close to 1 in the case of a strong positive correlation. We also report partial correlation coefficients and their posterior probabilities, which indicate the relationship between variables of interest after accounting for effects of all other covariates.

### Correlations between migration distance and molecular evolutionary rates (dS and dN/dS)

Our analyses with Coevol show that migration distance has a negative relationship with dS across the 39 species we studied, conforming to our initial predictions (Fig. 2, Fig. S2). For Coevol models with the full species set, the correlation coefficient between migration distance and dS was −0.39 with a posterior probability (*pp*) of 0.018, indicating strong support for a negative relationship. The partial correlation coefficient (which controls for the effects of mass) between migration distance and dS was −0.47 (*pp* = 0.0090).

We did not detect evidence of a relationship between migration distance and dN/dS (correlation coefficient = 0.096, *pp* = 0.63). The partial correlation coefficient (accounting for effects of mass) between migration distance and dN/dS showed weak support for a relationship (partial correlation coefficient = 0.26, *pp* = 0.82).

Results from the Coevol models of the subset of 30 species for which we had estimates of θ were consistent with results produced by the full subset (39 species) models, although support for the correlation between dS and migration distance was slightly weaker. In the model estimating dS, migration distance had a correlation coefficient of −0.43 (*pp* = 0.02) and a partial correlation coefficient of −0.31 (*pp* = 0.11). In the model estimating dN/dS, we did not find support for a relationship with migration distance, as this variable had a correlation coefficient of −0.15 (*pp* = 0.32) with dN/dS and a partial correlation coefficient of −0.010 (*pp* = 0.52) with dN/dS.

### Correlations between mass and molecular evolutionary rates (dS and dN/dS)

Our Coevol models with the full species set support the expected negative relationship between mass and dS (correlation coefficient = −0.28, *pp* = 0.065; Fig. 2). This relationship weakens when effects of migration distance are accounted for (i.e., with partial correlation coefficient = −0.18, *pp* = 0.20). We did not find a strong correlation between mass and dN/dS (correlation coefficient = −0.25, *pp* = 0.19; partial correlation coefficient = −0.072, *pp* = 0.41). In models of dS with the subset of 30 species that included θ as a predictor, mass had a correlation coefficient of −0.30 (*pp* = 0.10) and a partial correlation coefficient (which controls for the effects of migration distance) of −0.40 (*pp* = 0.039). In models of dN/dS from this subset, mass had a correlation coefficient of 0.25 (*pp* = 0.8) and a partial correlation coefficient of 0.23 (*pp* = 0.79).

### The influence of N_e_ on molecular rates and their correlation with traits of interest

In models using the subset of 30 species with population-level data, we did not find evidence for a correlation between θ and dS (correlation coefficient = −0.23, *pp* = 0.15; partial correlation coefficient = −0.12, *pp* = 0.67). This result is consistent with neutral evolution of synonymous sites among the species we studied. By contrast, we found strong support for the nearly neutral theory’s predicted negative relationship (Ohta 1992; Popadin et al. 2007; Leroy et al. 2021) between θ and dN/dS (correlation coefficient = −0.60, *pp* = 0.025; partial correlation coefficient = −0.57, *pp* = 0.031; Fig. 3), indicating stronger purifying selection in species with higher N_e_.

**Figure 3.**
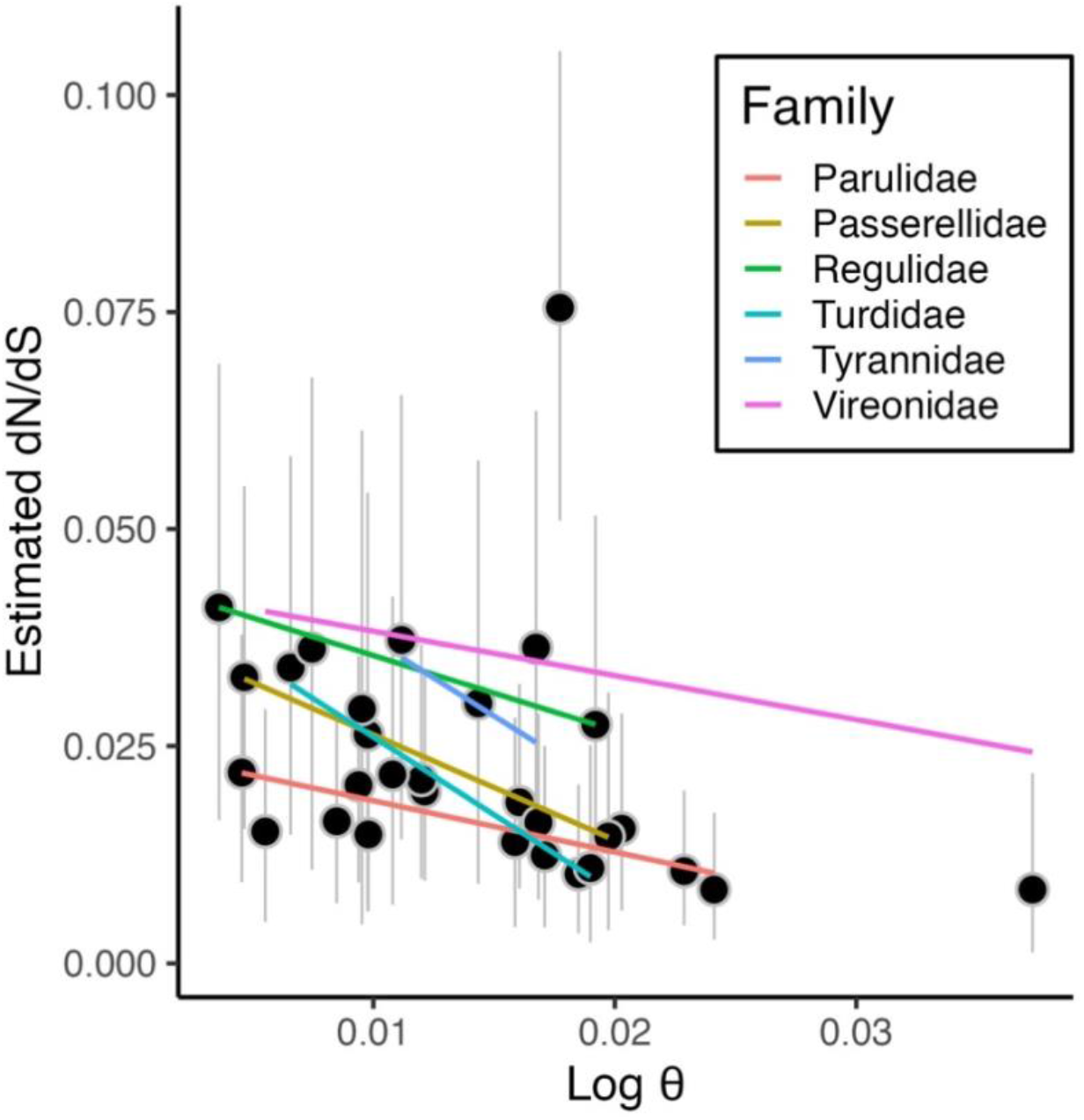
dN/dS vs. θ. Posterior mean tip estimates (black dots) of dN/dS are shown compared to θ from a Coevol model including species for which we could estimate θ. Gray vertical bars indicate 95% credible intervals for each estimate. As in Fig. 2, plotted lines use linear models to visualize the relationship between mean tip dN/dS and θ within each family of birds (when represented in our dataset by two or more species), demonstrating a consistently negative relationship between θ and dN/dS within and among major clades in our system.

### Linear modeling of πN/πS

In comparison of AICc, the highest-ranked model of πN/πS showed a strongly supported negative relationship between θ and πN/πS (Fig. 4, Table S6, model weight 0.55), as predicted if purifying selection is stronger in species with higher N_e_. Compared to a model with θ alone, a model with both θ and migration distance shows an increase in multiple r^2^ from 0.15 to 0.28 and a decrease in AICc by more than two units, suggesting the inclusion of migration distance improves model fit. However, contrary to our prediction, migration distance has a weak positive relationship with πN/πS (Fig. 4). The estimated coefficient relating θ and πN/πS in the best-fit model is −0.027 (std error = 0.01) and the estimated effect of migration distance from the best-fit model is 0.022 (std error = 0.01). Model comparison did not support the inclusion of mass as a predictor of πN/πS (Table S6).

**Figure 4.**
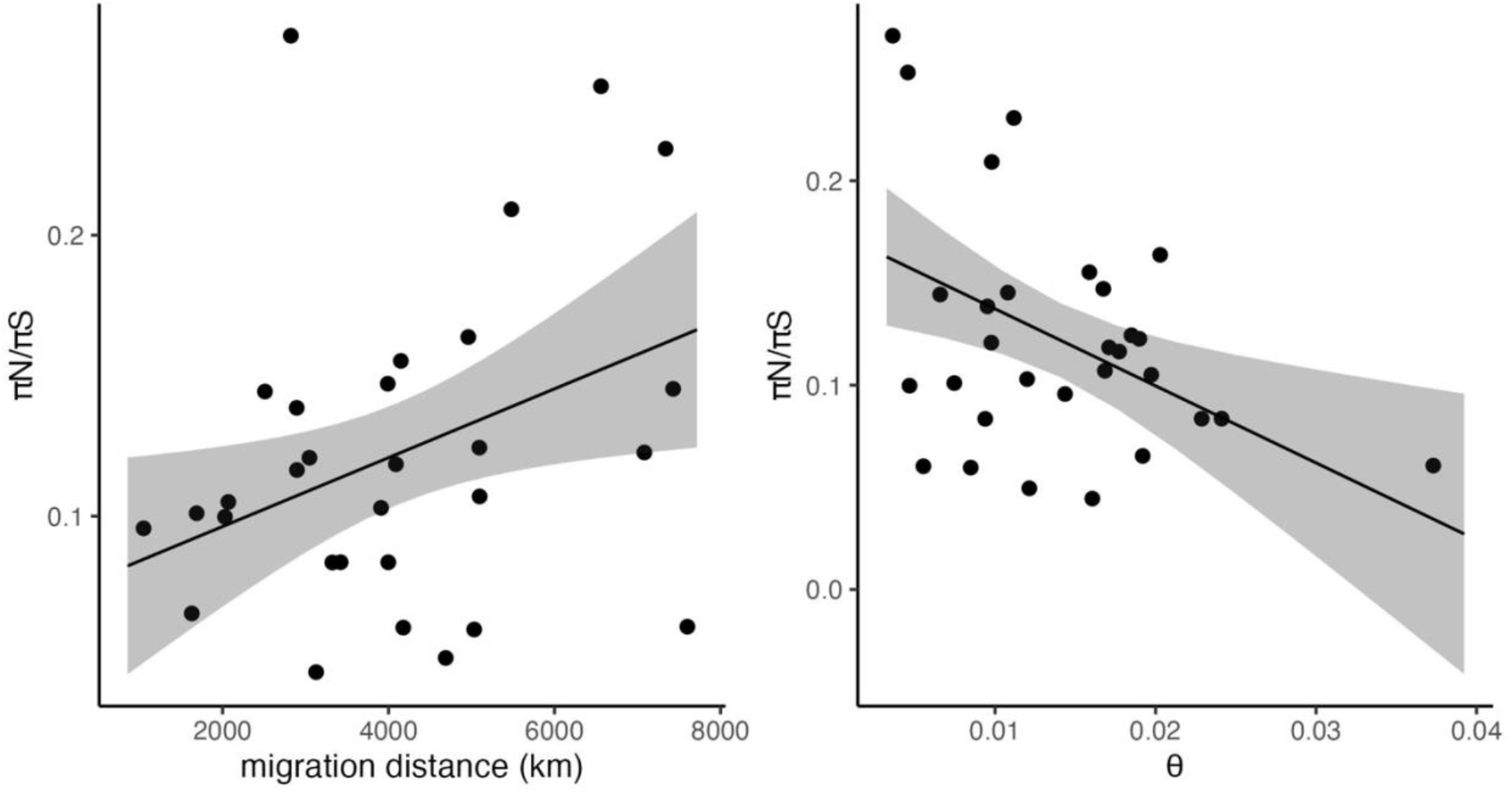
The relationship between πN/πS and migration distance (left) and θ (right). πN/πS is strongly influenced by θ, as expected if purifying selection removes more nonsynonymous variation in species with larger N_e_. πN/πS increases with migration distance, after accounting for effects of θ. Regression lines and 95% confidence intervals show the marginal effect of each variable as calculated by “ggpredict()” from the R package ggeffects v0.16.0 (Lüdecke 2018) using the best-fit model, which included both predictors.

### N_e_ is unlikely a confounding factor in inferred relationships

We used linear modeling to test whether migration distance or mass show a relationship with θ, our proxy of N_e_. We did not find strong evidence that body mass or migration distance are correlated with θ among the 30 species we studied. The null model for θ (an intercept-only model with no predictors) showed the lowest AICc, suggesting that the addition of mass and migration distance as predictors did not improve model fit (Table S7, model weight 0.45). However, the model with migration distance as a predictor was within 2 AICc units of the null model and showed a model weights of 0.30, indicating considerable model uncertainty. The estimated effect of migration distance on θ was positive but had a negligible effect size in the second-best model (estimate = 0.0017, std error = 0.0013 model multiple r^2^ = 0.054).

## Discussion

### Seasonal migration distance correlates with mitochondrial dS

We examined the relationship between life history and patterns of mitochondrial sequence evolution within North American boreal birds. These species occupy a region where strong seasonality demands specialized adaptations that carry life history tradeoffs (Varpe 2017; Winger and Pegan 2021). Our results implicate the life history axis of seasonal migration distance as a novel correlate of mitochondrial synonymous substitution rate (dS). Previous work demonstrates that, even after accounting for body size, long-distance migrants in this system have slower life history strategies than short-distance migrants, showing higher annual adult survival and lower fecundity (Winger and Pegan 2021). Here, we find that the slow life history of long-distance migrants is accompanied by a slower rate of neutral molecular evolution in the mitochondria of these species compared with that of shorter-migrating species in the region. Indeed, among the 39 species we studied, the correlation between migration distance and dS is stronger than the correlation between mass and dS, which is notable given that the relationship between mass and substitution rate has been documented in previous work (Nabholz et al. 2016). As such, we suggest that the association between migration distance and the slow-fast life history continuum extends to effects on dS.

### What evolutionary processes link migration distance with mitochondrial dS?

Substitution rates are fundamentally influenced by mutation rate, which provides new molecular variants with potential to become substitutions, and by natural selection, which influences whether variants are fixed as substitutions or lost. The correlation between migration distance and dS therefore reflects one or both processes. dS is often treated as a proxy for mutation rate alone based on the assumption that natural selection does not operate on synonymous sites (Nei et al. 2010), but in some cases synonymous sites are known to evolve non-neutrally (Chamary et al. 2006). If synonymous sites are not evolving neutrally, nearly neutral theory suggests that the relationship we find between dS and migration distance could hypothetically be explained by larger N_e_ in long-distance migrants (Ohta 1992). However, we did not find evidence for a correlation between dS and our proxy for N_e_ (θ) (Table S4) nor correlation between θ and migration distance (Table S7). Together, these results suggest that synonymous sites are evolving neutrally in our system and that variation in dS among species with different migration distances is not well explained by variation in demographic parameters that influence substitution rate. Rather, we suggest that the negative relationship we found between migration distance and dS may reflect a negative relationship between migration distance and mutation rate.

### Why might long-distance migrants have lower mitochondrial mutation rate?

We predicted that migration distance would correlate with dS because of its relationship with the slow-fast continuum of life history in these species independent of body size (Winger and Pegan 2021). In turn, a species’ position on the slow-fast life history continuum is hypothesized to affect mutation rate (Bromham 2020). There are several potential mechanisms to explain the link between life history and mutation rate, and the relative importance of each is not clear (Bromham 2020). The “copy error effect” hypothesis suggests that the explanation is related to generation time, assuming that “fast” species with short generation times and young age at first reproduction experience higher rates of germline replication (and thus replication-induced mutation) than species with “slow” life histories (Li et al. 1996; Thomas et al. 2010; Lehtonen and Lanfear 2014). However, recent studies comparing cell division rates with directly-measured mutation rates suggest that replication-induced copy errors may not be the only driver of differences in mutation rate between lineages (Wu et al. 2020; Wang et al. 2022). The “mutation avoidance” hypothesis offers another non-exclusive explanation for lower dS in organisms with slow life history based on higher costs of mutation in longer-lived species (Bromham 2020). Under this hypothesis, organisms with slow life history are predicted to have adaptations that reduce the introduction of mutations from DNA damage or from DNA replication and repair processes (Galtier et al. 2009; Tian et al. 2019; Zhang et al. 2021; Cagan et al. 2022). Long-distance migrants may be especially sensitive to costs of mitochondrial mutation, which may cause mitochondrial senescence (Galtier et al. 2009; Hua et al. 2015), because of the high physical performance demanded by their migratory behavior across their entire lifespans (Møller 2007; Conklin et al. 2017). Further research is necessary to understand what processes contribute to the apparent reduction of mutation rate in species at the slow end of the slow-fast continuum of life history.

Another possible link between migration distance and mutation rate is oxidative damage from metabolism, which is recognized as a potential source of mutation rate variation (Martin and Palumbi 1993, Gillooly et al. 2005, Berv and Field 2018; but see Lanfear et al. 2007, Galtier et al. 2009). Thus, a potential explanation for our results—lower mitochondrial dS in long-distance migrants—is that long-distance migrants incur less metabolically-induced DNA damage than do short-distance migrants. This explanation is initially surprising in light of studies showing that migratory birds experience oxidative damage from endurance flight (Jenni-Eiermann et al. 2014; Skrip and McWilliams 2016). However, we suggest that there are three plausible and non-exclusive scenarios that could lead to lower metabolically-induced DNA damage in long-distance compared to short-distance migrants. First, long-distance migrants may have better adaptations for flight efficiency (Weber 2009; Elowe et al. 2023), reducing the amount of oxidative damage they experience per mile traveled. Second, the mutation avoidance hypothesis predicts that long-distance migrants may have more efficient DNA repair mechanisms than short-distance migrants, which could reduce metabolically-induced mutation rate even when long-distance flight does induce high oxidative stress. Last, short-distance migrants in our boreal study system may experience greater oxidative damage arising from their increased need for winter cold tolerance than long-distance migrants that winter in the tropics.

The mitochondria also play an important role in the metabolic challenge of maintaining homeostasis during cold weather and resource shortages (Bicudo et al. 2001; Chen et al. 2018). Short-distance boreal migrants likely face more of these kinds of challenges than long-distance migrants during both migration and winter (Winger and Pegan 2021). Despite the view that long-distance migration is an extreme performance challenge, its alternative—spending the winter within the temperate zone—is also an extreme metabolic challenge for small-bodied homoeothermic endotherms that do not hibernate (Dawson and Yacoe 1983; Winger et al. 2019). Further investigation of the comparative metabolic challenges faced by short versus long distance boreal migrants is needed to clarify whether and how migration distance influences metabolically-induced mutation in the mitochondria.

### Purifying selection is not stronger in long-distance migrants

Whereas evolutionary rate at synonymous sites (dS) may primarily reflect mutation rate, evolution at nonsynonymous sites is expected to strongly reflect natural selection because nonsynonymous mutations alter the amino acid sequence of a gene’s protein product. We found that the ratio of nonsynonymous to synonymous substitutions (dN/dS) among our species is universally much less than 1 (Fig. 3), indicating that the mitochondrial genes we studied are under purifying selection in all species in the system. We similarly found low ratios of nonsynonymous to synonymous polymorphisms within each population (πN/πS; Fig. 4), which is also consistent with purifying selection. Moreover, both dN/dS and the πN/πS ratio are strongly correlated with θ, our proxy for N_e_ (Fig. 3, 4), as expected under nearly neutral theory (Ohta 1992). A complexity of our results is that dN/dS reflects the accumulation of substitutions across the entire history of a lineage, whereas population parameters such as θ and πN/πS may be more strongly influenced by recent demographic processes. However, that we and others (e.g., Popadin et al. 2007; Leroy et al. 2021) find empirical evidence for the relationship between θ and dN/dS predicted by nearly-neutral theory, despite this potential mismatch in evolutionary timescales, suggests that empirical estimates of genetic diversity and molecular evolutionary rates may be shaped by similar demographic processes.

Our results are consistent with the general finding that mitochondrial genes tend to experience strong purifying selection (Nabholz et al. 2013; Popadin et al. 2013). However, we did not find evidence supporting our prediction that long-distance migrants would show stronger purifying selection (i.e., lower dN/dS and πN/πS) than short-distance migrants. This finding may reflect the fact that all species in our system face generally strong mitochondrial purifying selection, such that the endurance flights of long-distance migrants do not incur much stronger selection than the level that exists among all the species we studied. Our results also imply that short-distance migrants in the boreal region do not experience *relaxed* purifying selection on mitochondrial genes compared to long-distance migrants. As noted above, short-distance boreal migrants contend with metabolic challenges associated with cold winter temperatures in addition to the metabolic demands of flight, which may also exert selection on the mitochondria (Chen et al. 2018).

### Migration distance and the costs of mitochondrial mutations

In this study, we based our predictions on several complementary hypotheses about the costs of mutation in species with slow life history and high demand for physiological performance, such as long-distance migrants. From the perspective of molecular evolution, the mutation avoidance hypothesis (Bromham 2020) and studies on the relationship between lifespan and mutation rate (Nabholz et al. 2008a; Galtier et al. 2009; Tian et al. 2019; Zhang et al. 2021) predict that phenotype-altering genetic variation is harmful enough to induce selection for mutation avoidance in organisms with slow life history. From the perspective of population biology, the hypothesis proposed by Conklin et al. (2017) predicts that “slow” species with high performance demands experience a strong selective filter on phenotypic performance in early life, reducing phenotypic variation in these populations. While Conklin et al. (2017) frame their hypothesis around reduction of phenotypic variation, a similar prediction about reduction of genetic variation emerges from a series of studies showing that mitochondrial purifying selection is stronger in species with higher locomotory metabolic demands (Shen et al. 2009; Chong and Mueller 2013; Strohm et al. 2015; Mitterboeck et al. 2017; Sun et al. 2017; Chang et al. 2020; De Panis et al. 2021). Together, these hypotheses led us to predict that costs of mitochondrial mutation in long-distance migrants, which have slow life history, would cause them to exhibit slower mitochondrial mutation rate and stronger mitochondrial purifying selection than short-distance migrants.

Our predictions were only partially supported. The negative relationship we found between migration distance and dS is consistent with lower mitochondrial mutation rate in long-distance migrants, but we did not find evidence that these species experience stronger mitochondrial purifying selection than do short-distance migrants. To reconcile these findings and advance our understanding of how long-distance migration influences molecular evolutionary dynamics, further research is needed on the relative metabolic demands of long-distance flight versus cold tolerance and on the consequences of mitochondrial genetic variation for migratory phenotype. Additionally, studying molecular rates across the nuclear genome will also help clarify which dynamics we report here are related to selection on the mitochondrial genome and which reflect more general interactions between life history and molecular evolution.

### Conclusions: seasonal adaptation provides novel context for studying the links between life history and molecular evolutionary rates

Adaptation to seasonality entails life history tradeoffs (Varpe 2017). Organisms balance these tradeoffs in different ways, creating variation in life history strategy within communities that inhabit seasonal environments (e.g., Winger and Pegan 2021). Our study demonstrates that life history variation related to seasonality can influence molecular evolutionary rate, which has potential implications for accurate reconstruction of evolutionary history (Shafir et al. 2020; Ritchie et al. 2022). More broadly, we suggest that communities adapted to seasonal habitats provide an interesting context in which to investigate potential drivers of the relationship between life history and molecular evolution. Co-distributed species show varying adaptations to seasonality—e.g., cold tolerance, migration, hibernation—and they express these strategies to different degrees (Auteri 2022). Cold adaptations can influence biological processes hypothesized to be relevant for germline replication rate or mutation rate (e.g., Wang et al. 2022), even among species that show little variation in commonly-studied life history proxies such as body mass. Comparative studies using seasonal communities can therefore allow us to draw new insights into how life history tradeoffs affect mutation rate, one of the most fundamental processes in evolution.

## Acknowledgements

We thank Natalie Hofmeister, Kristen Wacker, Matt Hack, Susanna Campbell, Andrea Benavides Castaño, and the lab of Stephen Smith for helpful discussion. Teia Schweizer, Christine Rayne, and Kristen Ruegg provided lab assistance. For field sampling permits, we thank the United States Fish and Wildlife Service, the United States Forest Service, the Minnesota Department of Natural Resources, the Michigan Department of Natural Resources, the Canadian Wildlife Service of Environment and Climate Change Canada, Alberta Fish and Wildlife, and Manitoba Fish and Wildlife. Field sampling was approved by the University of Michigan Animal Care and Use Committee. For providing additional samples, we thank the American Museum of Natural History (Brian Smith, Joel Cracraft, Paul Sweet, Peter Capainolo, Tom Trombone), Royal Alberta Museum (Jocelyn Hudon), University of California, Berkeley Museum of Vertebrate Zoology (Rauri Bowie and Carla Cicero), Cleveland Museum of Natural History (Andrew Jones, Courtney Brennan), Cornell University Museum of Vertebrates (Irby Lovette, Vanya Rohwer, Mary Margaret Ferraro, Charles Dardia), University of Michigan Museum of Zoology (Brett Benz, Janet Hinshaw), University of Minnesota Bell Museum of Natural History (Keith Barker), and the New York State Museum (Jeremy Kirchman). For assistance in the field, we thank Brett Benz, Courtney Brennan, Susanna Campbell, Shane DuBay, Ethan Gyllenhaal, Mary Margaret Ferraro, Laura Gooch, Andrew Jones, Heather Skeen, Vera Ting, and Brian Weeks. Next-generation sequencing for this project was partially carried out in the Advanced Genomics Core at the University of Michigan. This research was also supported in part through computational resources and services provided by Advanced Research Computing (ARC), a division of Information and Technology Services (ITS) at the University of Michigan, Ann Arbor. This material is based upon work supported by the National Science Foundation under Grant No. 2146950 to BMW. This research was supported by the Jean Wright Cohn Endowment Fund, Robert W. Storer Endowment Fund, Mary Rhoda Swales Museum of Zoology Research Fund and William G. Fargo Fund at the University of Michigan Museum of Zoology. TMP was supported by the NSF Graduate Research Fellowship (DGE 1256260, Fellow ID 2018240490) and a University of Michigan Rackham Graduate Student Research Grant.

---

**Figure S1.**
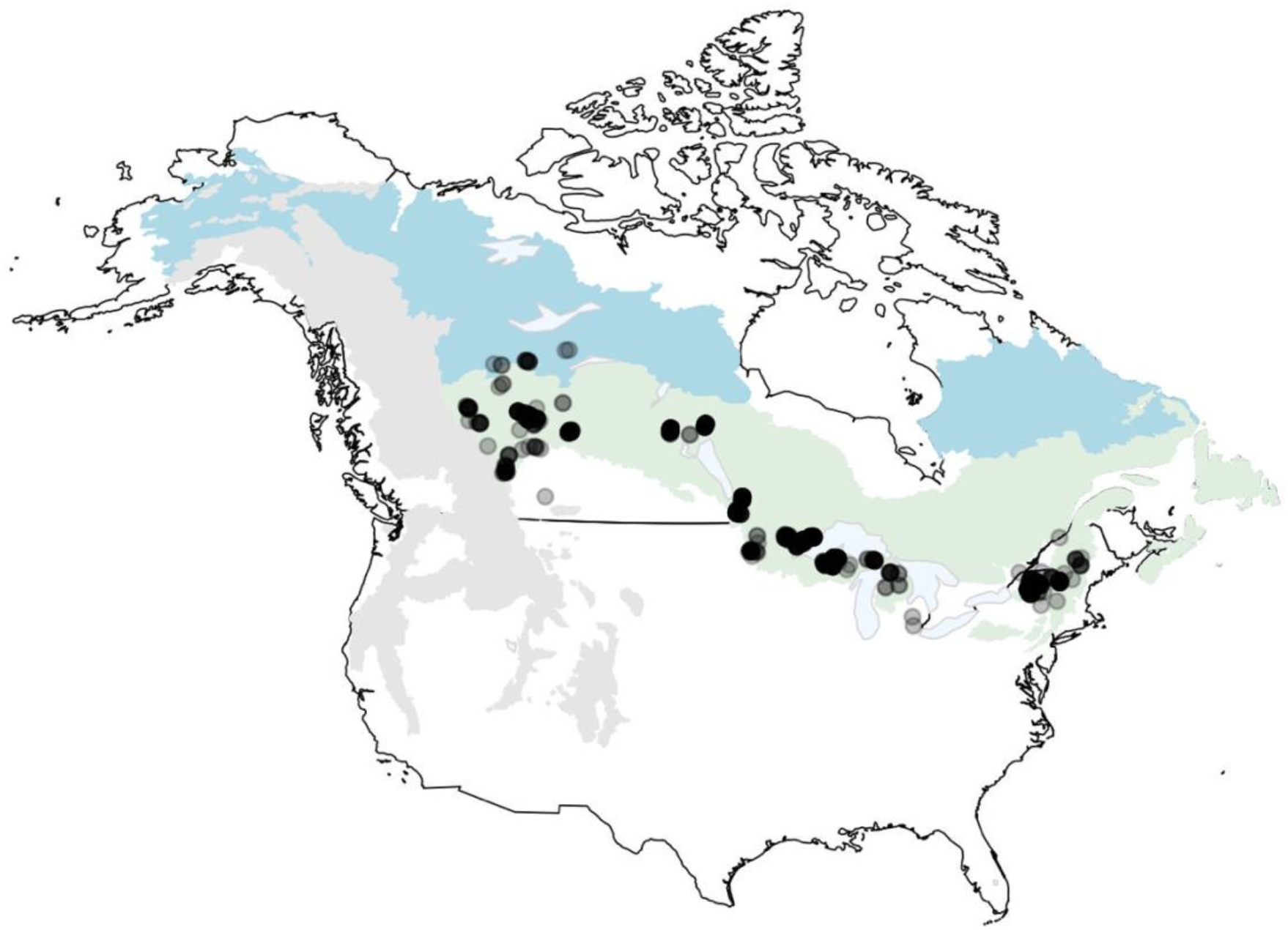
Map of sampling locations for all 39 species. Each point represents an individual, such that darker shading indicates multiple individuals. The boreal forest (green), taiga (blue) and Rocky Mountains (grey) are designated following ‘level 1’ ecoregions defined by Omernik and Griffith (2014).

**Figure S2.**
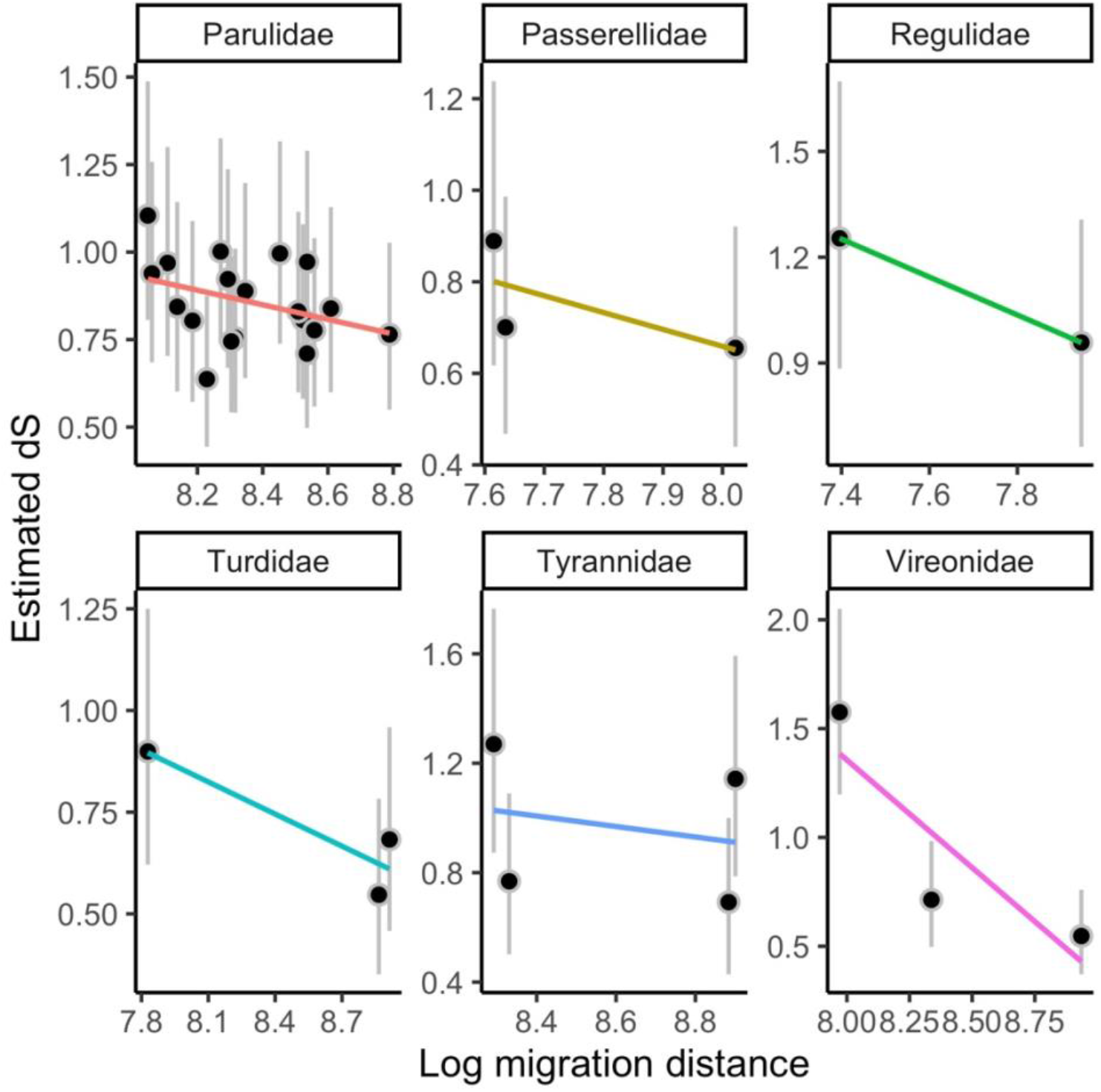
The relationship between dS and migration distance within each family represented in our study by more than one species. Posterior mean tip estimates of dS (black dots) from Coevol are shown compared to migration distance (left), and mass (right) from models using our full species set. Gray vertical bars indicate 95% credible intervals for each estimate. Plotted lines use linear models to visualize the relationship between estimated tip dS and a given covariate within each family, demonstrating a consistently negative relationship between dS and migration distance within and among major clades in our system.

**Table S1.** [provided as a separate spreadsheet]. Species used in this study with information about each sequence used in Coevol analyses (sample catalog number); previously-published GenBank sequences used during sample processing for each species (seed sequence provided to NOVOPlasty and mitochondrial coding sequences used for annotation with Geneious); data used in this study (mass and migration distance from sources described in the methods, θ as calculated in this study); the number of samples used to calculate population genetic summary statistics; posterior mean estimates of dS and dN/dS with upper and lower 95% credible intervals; and piN/piS estimates. Estimates of dS and dN/dS (and credible intervals) each come from one replicate of a Coevol model using the entire 39-species dataset and serve as representative estimates. Detailed information about each sample, including all of those used in population genetic analyses, can be found in Table S2.

**Table S2.** [provided as a separate spreadsheet]. Information about each sample used in this study, including its catalog number and institution, sex, date and locality of collection (state or province within the US or Canada and geographic coordinates), whether the sample was ultimately removed (retained = FALSE) from analysis because of low-quality data or other problems (1008 out of 1229 samples were retained), and the GenBank accession number for each mitochondrial gene. Museum abbreviations: AMNH = American Museum of Natural History; CMNH = Cleveland Museum of Natural History; CUMV = Cornell University Museum of Vertebrates; MMNH = Bell Museum of Natural History; MVZ = Museum of Vertebrate Zoology, UC Berkeley; NYSM = New York State Museum; RAM = Royal Alberta Museum; UMMZ = University of Michigan Museum of Zoology.

**Table S3.** Coevol models using alternative topologies produce consistent results. This table summarizes Coevol’s estimated relationship between dS and migration distance across models using 10 different marginal trees sampled from our phylogenetic tree dataset (see Methods). We show correlation coefficients and partial correlation coefficients and their associated posterior probabilities. Despite variation in topology across the marginal trees, these models each estimate similar correlation coefficients and similar levels support from posterior probabilities.

| **Sampled tree** | **Cor. Coeff.** | **Cor. Coeff. Post. Prob.** | **Part. Cor. Coeff.** | **Part. Cor. Coeff. Post. Prob.** |
| --- | --- | --- | --- | --- |
| 1 | -0.348 | 0.046 | -0.379 | 0.03 |
| 2 | -0.355 | 0.046 | -0.405 | 0.023 |
| 3 | -0.331 | 0.054 | -0.371 | 0.029 |
| 4 | -0.326 | 0.058 | -0.303 | 0.067 |
| 5 | -0.275 | 0.094 | -0.274 | 0.089 |
| 6 | -0.349 | 0.047 | -0.379 | 0.029 |
| 7 | -0.305 | 0.067 | -0.297 | 0.072 |
| 8 | -0.285 | 0.081 | -0.28 | 0.079 |
| 9 | -0.325 | 0.06 | -0.321 | 0.057 |
| 10 | -0.351 | 0.045 | -0.406 | 0.021 |

**Table S4.** [provided as a separate spreadsheet]. Full output from Coevol models with dS and dN as independent variables. Four models are summarized: two replicate models with the full dataset of 39 species, and two replicate models with the subset of 30 species with available estimates of θ. Each row contains a flattened pairwise matrix for each pair of variables (dS, dN, mass, migration distance, and θ), showing covariance, correlation coefficients, posterior probabilities of correlation coefficients, precisions, partial correlation coefficients, and posterior probabilities of partial correlation coefficients. Posterior probabilities near 0 indicate strong support for a negative relationship while posterior probabilities near 1 indicate strong support for a positive relationship. Values obtained from replicate models are highly similar.

**Table S5.** [provided as a separate spreadsheet]. Full output from Coevol models with dS and dN/dS as independent variables. Four models are summarized: two replicate models with the full dataset of 39 species, and two replicate models with the subset of 30 species with available estimates of θ. Each row contains a flattened pairwise matrix for each pair of variables (dS, dN/dS, mass, migration distance, and θ), showing covariance, correlation coefficients, posterior probabilities of correlation coefficients, precisions, partial correlation coefficients, and posterior probabilities of partial correlation coefficients. Posterior probabilities near 0 indicate strong support for a negative relationship while posterior probabilities near 1 indicate strong support for a positive relationship. Values obtained from replicate models are highly similar.

**Table S6.** Full model selection results for models predicting πN/πS.

| Model formula | logLik | AICc | deltaAICc | weight |
| --- | --- | --- | --- | --- |
| $\pi N/\pi S \sim \theta + \text{migration distance}$ | 48.55 | -87.5 | 0 | 0.55 |
| $\pi N/\pi S \sim \theta + \text{mass} + \text{migration distance}$ | 48.68 | -84.9 | 2.63 | 0.15 |
| $\pi N/\pi S \sim \theta$ | 45.88 | -84.8 | 2.65 | 0.15 |
| $\pi N/\pi S \sim 1$ (null) | 43.53 | -82.6 | 4.89 | 0.048 |
| $\pi N/\pi S \sim \text{migration distance}$ | 44.70 | -82.5 | 5.02 | 0.045 |
| $\pi N/\pi S \sim \theta + \text{mass}$ | 45.88 | -82.2 | 5.33 | 0.038 |
| $\pi N/\pi S \sim \text{mass}$ | 43.55 | -80.2 | 7.31 | 0.014 |
| $\pi N/\pi S \sim \text{mass} + \text{migration distance}$ | 44.70 | -79.8 | 7.69 | 0.012 |

**Table S7.** Full model selection results for models predicting θ.

| Model formula | logLik | AICc | deltaAICc | weight |
| --- | --- | --- | --- | --- |
| $\theta \sim 1$ (null model) | 105.96 | -207.5 | 0 | 0.45 |
| $\theta \sim$ migration distance | 106.88 | -206.6 | 0.82 | 0.30 |
| $\theta \sim$ mass | 106.08 | -205.2 | 2.24 | 0.15 |
| $\theta \sim$ mass + migration distance | 107.10 | -204.6 | 2.88 | 0.11 |

## Notes

### Competing Interest Statement

The authors have declared no competing interest.

